# *Arbuscular Mycorrhizal Fungus* (AMF) and reduction of arsenic uptake in lentil crops

**DOI:** 10.1101/522714

**Authors:** Mohammad Zahangeer Alam, Md. Anamul Hoque, Rebecca McGee, Lynne Carpenter-Boggs

## Abstract

Arsenic (As) is a carcinogenic and hazardous substance that poses a serious risk to human health. Physiological studies have shown that growth of lentil crop have been impaired due to arsenic toxicity, and is transportable into human food chains. Our research focused on the transportation of As in lentil crops and its mitigation using Arbuscular Mycorrhizal Fungus (AMF). Shoot length, fresh and dry weight of shoot and root were found comparatively higher in 5 and 15 mgkg^-1^ arsenic treated lentil seedlings than in a 100 mgkg^-1^ As concentrated soil. As accumulation in lentil’s pods of BARI Mashur 1 were found higher than others; but As uptake in root and shoot were increased significantly in all BARI released lentil genotypes. Biomass growth of lentil was found higher in AMF treated soils in compare to non-AMF. AMF effectively reduced the arsenic uptake in root and shoot at 8 and 45 mgkg^-1^ As concentrated soils compared. As free lentil seeds are significantly important for human consumption through mitigation of As accumulation in lentil roots shoots and pods. AMF shows great potential in providing As free lentil seeds throughout the world.

## 1. INTRODUCTION

Arsenic (As) is a natural but hazardous element present in rocks, soils, water, air, and biological tissues (Hossain, 2006). Research has increased in recent years on the occurrence, distribution, origin, and mobility of As in soils through natural, geochemical and biological processes (Leung et al., 2013). According to the U.S. Agency for Toxic Substances and Disease Registry (ATSDR) priority list of hazardous substances, As has been designated as the number one hazardous substance in the United States (Leung et al., 2013). Moreover, As contamination has been reported worldwide in Argentina, Australia, Bangladesh, Chile, China, Hungary, Mexico, Peru, Thailand, and Vietnam (Ahmed et al., 2011). However, the most severe As contamination to surface soil, water, and humans is currently in Asia, particularly Bangladesh, (Ahmed et al., 2011) West Bengal and India (Ahmed et al., 2006).

Arsenic has been recognized as a carcinogenic substance based on its chemical and physical forms as well as concentration and duration of exposure (Singh et al., 2015). Chemically, it exists as organic and inorganic species. The main sources of arsenic are arsenic sulphide (As_2_S_2_), arsenic tri-sulphide (As_2_S_3_) and arsenopyrite or ferrous arsenic sulphide (FeAsS_2_) (Hossain, 2006). Inorganic As has two main oxidation states (i.e., trivalent arsenite As(III), and pentavalent arsenate As(V). The inorganic forms of arsenate As(V) and arsenite As(III)) are usually dominant in As contaminated soil. The arsenite As(III) in the presence of herbicides and pesticides is oxidized into As(V) (Cubadda et al., 2010). Trivalent arsenite is 60 times more toxic than arsenate (Hossain, 2006).

Arsenic causes highly toxic effects on metabolic processes of plants, mitotic abnormalities, leaf chlorosis, growth inhibition, reduced photosynthesis, DNA replication, and inhibition of enzymatic activities (Nagajyoti et al., 2010). For instance, root and leaf elongation of the mesquite plant *(Prosopis juliflora x P. velutina*) decreased significantly with increasing As (III) and As (V) concentrations (Ntebogeng et al., 2008). Heikens et al. (2006) reports that As contaminated water leads to accumulation in the soil, which is then transported into edible parts of food crops. Arsenite As(III) and arsenate As(V) both are present in wheat crops due to accumulation from soils to shoots and grains (Cubadda et al., 2010). In addition, the extensive use of pesticides, fertilizer, groundwater and industrial wastewater for irrigation purposes in crop fields has resulted in elevated levels of As in soils, and thus increased As uptake in rice, lentil and vegetables (Ahmed et al., 2011). Consequently, many food crops have become hazardous including Lentil, which is a major leguminous crop across the world. These crops are an excellent source of protein, minerals and vitamins for human nutrition (Guillon and Champ, 2002). Similarly, chronic exposure of As has led to unacceptable As levels in samples of soils, water, vegetables and cereals. Subsequently, high Average Daily Dose (ADD) from the environment and low excretion could result in As toxicity to humans from lentil crops as well as from other food crop cultivation in As contaminated soils (Cui et al., 2013). Furthermore, As carcinogenicity has caused serious health diseases, such as lung and skin cancers, and possible damage to liver and kidneys as well. Noncancerous health effects of As exposure include diabetes, chronic cough, and cardiovascular and nervous system collapse (SOS, 2011).

Currently, Bangladesh is the second largest area of As contamination in the world. Bangladesh is facing a serious public health threat, with 85 million people at risk of As contamination in drinking water and food crops. In addition, 85–95% of rice, lentil and vegetable crops are contaminated by As, which poses a serious threat to human and livestock health (Hossain, 2006). Therefore, it is imperative for the mitigation of As in crop plants. One possible solution includes Arbuscular Mycorrhizal Fungi (AMF), which establishes a mutualistic symbiotic relationship with the majority of terrestrial plant including lentil crops (Schneider et al., 2013). AMF are actively involved in As accumulation, and affects the concentration of As, Cd, Zn, and Pb in shoots and roots (Orloska et al., 2012). The effect of AMF on element uptake can, vary largely, depending on plant species/cultivar and metal concentration in the soil, but also on AM fungal species and isolates (Orloska et al., 2012). In aerobic soils the main form of As is arsenate As(V). In this form As mimics phosphorus (P), and can be taken up by lentil plants and AMF by normal P uptake mechanisms (Toulouze et al., 2012). In this circumstance, mycorrhizal symbioses are significantly highlighted because they are formed by 90% of higher plants, often with increased uptake of phosphate (P) compared with non-mycorrhizal (NM) counterparts (Smith et al., 2010). It is clear that the association of AMF inoculation with lentil crops might reduce As uptake by various mechanisms (Ahmed et al., 2011). The high proportion of inorganic species of As (As*i*) is of particular concern to the human carcinogen through the protein sources of lentil crops. Lentil is one of the major leguminous crops in the world. The future of agriculture will depend increasingly upon legume crops because of production of high energy and protein for human and animal health nutrition. Therefore, As mitigation technique is very much a necessity for lentil crops as well as other crops. The present research focused to lentil varietal selection against As and its impact on lentil’s biomass. This research also highlights the reduction of As accumulation in roots, shoots and pods using the Arbuscular Mycorrhizal Fungus (AMF). It hypothesized that this research is significantly important for the exploration of high and low As accumulator lentil that will supply arsenic free pods for the consumption to human populations.

## 2. MATERIALS AND METHODS

### 2.1. Arsenic accumulation in lentil roots and shoots

#### Soil sampling areas

Arsenic contaminated soils were collected for this pot experiment from Mathchar, Bangladesh Jute Research Institute (BJRI) area (Faridpur) and Bangabandhu Sheikh Mujibur Rahman Agricultural University (BSMRAU) research field (Gazipur) of Bangladesh, 2015. The Global Positioning System (GPS) are 23°35.38969’, 24°2.17859’, & 23°35.97636’ Latitudes and 89°48.69921’, 90°23.83393’, & 89°46.7586’ Longitudes in the soil sampling locations of BJRI, BSMRAU and Mathchar, respectively.

#### Collection of lentil genotypes

Bangladesh Agricultural Research Institute (BARI) is developed eight lentil varieties. Among these, 7 lentil varieties were procured from BARI for this study. These lentil varieties are BARI Mashur1, BARI Mashur 2, BARI Mashur 3, BARI Mashur 4, BARI Mashur 5, BARI Mashur 6 and BARI Mashur 7 (Table 1).

**Table 1.**
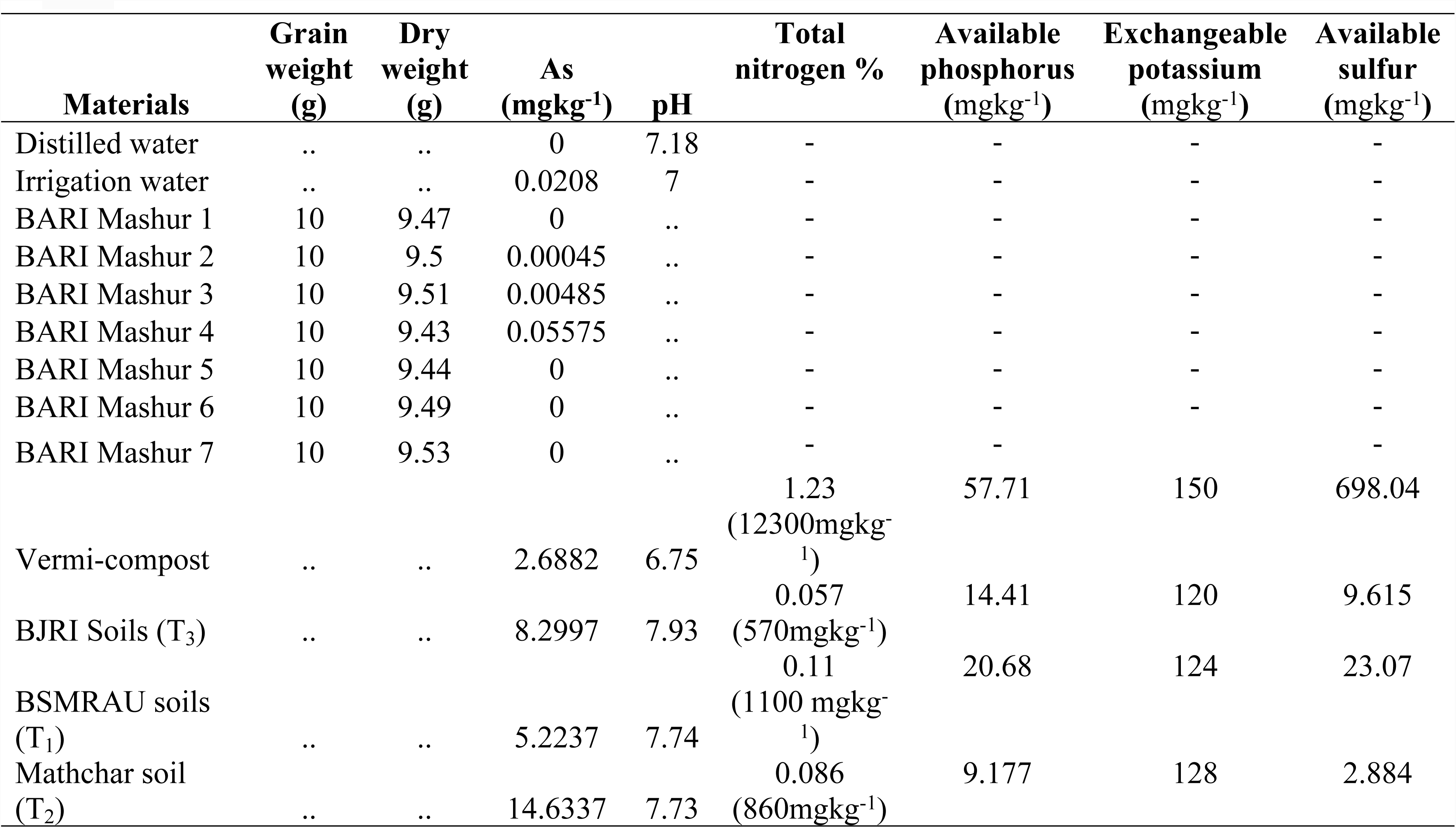
Dry weight and chemical properties of lentil seeds, soil, and water samples

#### Collection of vermi-compost, mineral fertilizers, brick’s pots and fungicides

Vermi-compost mixed with soils equally in all treated pots. Urea, Triple Super Phosphate (TSP) and Muriate of Potash (MOP) applied in soils of this experiment as source of Nitrogen (N), Phosphorus (P) and Potassium (K), respectively. Vitavex 200 fungicides used as seed treating chemical for lentil seeds. Clay pots size 6″/6″were used in this experiment. All types of input materials purchased from the local market of Bangladesh for this pot experiment.

#### Samples preparation

Soil samples collected from As contaminated regions in Bangladesh using a soil auger to a depth of 15 cm and brought into the Department of Environmental Science at Bangabandhu Sheikh Mujibur Rahman Agricultural University (BSMRAU). Before sowing Lentil seeds in a pot, initial soil samples of 250-300 (g) were taken from each composite through the guidelines of BARC (2012). The soil was air dried at room temperature in the laboratory. Samples were then ground and sieved with a ≤250 µm mesh and preserved in polythene bags with proper labeling. Vermi-compost samples were also prepared for chemical analysis as well as soil samples. Similarly, seeds, roots and shoots of lentil genotypes were kept in an oven for drying at 55°C for 72 hours. Samples were then ground using coffee grinder and liquid nitrogen and sieved with ≤250 µm mesh.

#### Chemical properties of soil, water, vermi-compost and dry weight of lentil seeds

The pH of soil, irrigation water and vermicompost were determined by glass electrode pH meter (Jackson 1973). Total N percentage of soil and vermi-compost were determined by Kjeldhal systems (Jackson, 1973). Available P of soil and vermi-compost were determined by Olsen, the Bray and Kurz method (Olsen and Sommers, 1982). Exchangeable K of soil and vermi-compost were estimated by Ammonium Acetate Extraction method (Jackson, 1973). Available sulfur of soil and vermi-compost were determined by turbidimetrically as barium sulfate method (Chesnin and Yien, 1951). Dry weight of lentil seeds was measured by digital electrical balance (Table 1).

#### Mixing of soil, vermi-compost and fertilizers substrate in pot

Collected soil samples were ground uniformly for sowing of lentil seeds. 1500 g ground soils with 200g vermi-compost were mixed in each pot. According to the recommendation of Bangladesh Agricultural Research Institute (BARI), Urea 225kgha^-1^, TSP 450kgha^-1^ and MOP 175 kgha^-1^ were incorporated with soils in each pots. Total nitrogen (61.33 mgkg^-1^), phosphorus (56.66 mgkg^-1^) and potassium (66.66 mgkg^-1^) were added, in each experimental pot from synthetic fertilizers. Then 7-10 lentil seeds of each variety sowed in each pot during the first week of November in 2015.

#### Treatments and replications in pot experiment

Based on the analysis of total arsenic content in soil samples, three soil samples were selected for treatments. These treatments included T_1_ = total arsenic content 5 mgkg^-1^ (BSMRAU soil), T_2_ = total arsenic content 15 mgkg^-1^ (Mathchar soil-Faridpur) and T_3_ = total arsenic content (8+92) =100 mgkg^-1^ (BJRI soil-Faridpur). Five replications with seven lentil varieties were used in these experiments with a 105 total number of pots.

#### Average shoot length, fresh and dry weight of lentil seedlings

At random, average shoot lengths were measured using a measuring tape (cm) at week 3 in each treated pots. At this time point, three lentil seedlings were thinned out from each arsenic treated pot. Fresh weights were taken of each sample using electrical balance (g). Average dry weight of roots and shoots were measured separately after harvesting of lentil seedlings from each As treated pot during week nine. All samples were dried in an oven at 55°C for 72 hours towards the digestion of samples for the determination of total As accumulation in root and shoot of lentil crops from soil samples.

### 2.2. Arsenic uptake in lentil pods during field condition

Simultaneously, **s**even lentil genotypes were sown on 12 November 2015 in field soils. For this field experiment, 10 x 5-meter sizes of seven plots were prepared at BSMRAU research fields. BARI released seven genotypes sown in seven plots separately. All plots were 5 mgkg^-1^ As concentrate soils. Recommended doses of fertilizers were applied to previous pot experiments. Lentil seedling harvested on 16 February 2016. Total duration was required 95 days from sowing to harvesting time of lentil crops. During harvesting, three samples of lentil pods were randomly collected separately from each plot and tagged with proper marking of each sample. Then samples were dried at room temperature. Next, all samples were dried in an oven at 55°C for 72 hours towards the digestion of samples for the determination of total As accumulation in lentil’s pods from soil samples.

### 2.3. Mitigation of arsenic through mycorrhizal inoculation

#### Selection of lentil genotypes

Based on the pervious field experiments, BARI Mashur 1 and BARI Mashur 5 were selected for the mitigation of As uptake through mycorrhizal inoculation. These pot experiments were conducted in a green house with a controlled environment at BSMRAU.

#### Collection of Arbuscular Mycorrhizal Fungus (AMF)

AMF samples were collected from International Culture Collection of (Vesicular) Arbuscular Mycorrhizal Fungi (INVAM), West Virginia University (WV), USA. AMF samples were mixed with soils and roots of the host plant of Sorghum that was housed in the Department of Environmental Science at BSMRAU. Mixture of soil and roots were collected from this cultured area as a source of AMF. Finally, this mixture of AMF was used for the reduction of As uptake in lentil roots, and shoots.

#### Observation of mycorrhizal spores and root colonization

Mycorrhizal spores in soil were extracted by following the Wet Sieving and Decanting Method (Gerdemann and Nicolson, 1963). Soil samples were collected from rhizosphere of Sorghum and mixed thoroughly. Unwanted particles such as stones, roots, and twigs were removed from these samples as needed. From this mixture, 100 g of soil samples were kept in a bucket with three quarters of tap water (~8 Liter). This mixture was stirred vigorously by hand and washed into a bucket and left to settle for one minute. This suspension was sieved by 400 μm and 200μm mesh throughout the experiment. Next, collected samples were poured through a 100μm sieve into a second bucket (10 litters) to avoid the loss of useful materials. After suspension settled for one minute, the supernatant was decanted using a 400-μm sieve and the water was discarded. The solution with spores was distributed into 4 equal size test tubes using water for equal weight. The tubes were plugged properly and then centrifuged for 4 minutes at 3,000 rpm. The supernatant was then poured in the test tubes, filled with sucrose solution, and stirred vigorously with the round-ended spatula re-suspended precipitate. The plugged test tubes were then centrifuged for 15 seconds at 3,000 rpm. After centrifugation, the sucrose supernatant was poured through a 400μm sieve and rapidly washed with water to remove the sucrose from AMF spores by back washing the materials from the sieve into a wash glass for observation. The spores in the wash glass were observed under Stereomicroscope and transferred to microscope slides. Then the slide placed under an electron-microscope for the observation of their size. **Similarly**, the sorghum root was rinsed thoroughly in water and cut into small pieces, then placed in 2.5% KOH solution. Roots were then heated in a water bath at 90°c for 10-30 minutes and kept in 1% HCl solution overnight. Then samples were stained in acidic glycerol with 0.05% aniline blue for 10-30 minutes at 90°c. The de-stained samples were left at room temperature in acidic glycerol. Similarly, the roots were kept on the slides and observed under an electron microscope for the observation of spores’ size, and its attachment with mycelia and hyphae (Giovanetti and Mosse, 1980) (Figure 1).

**Figure 1.**
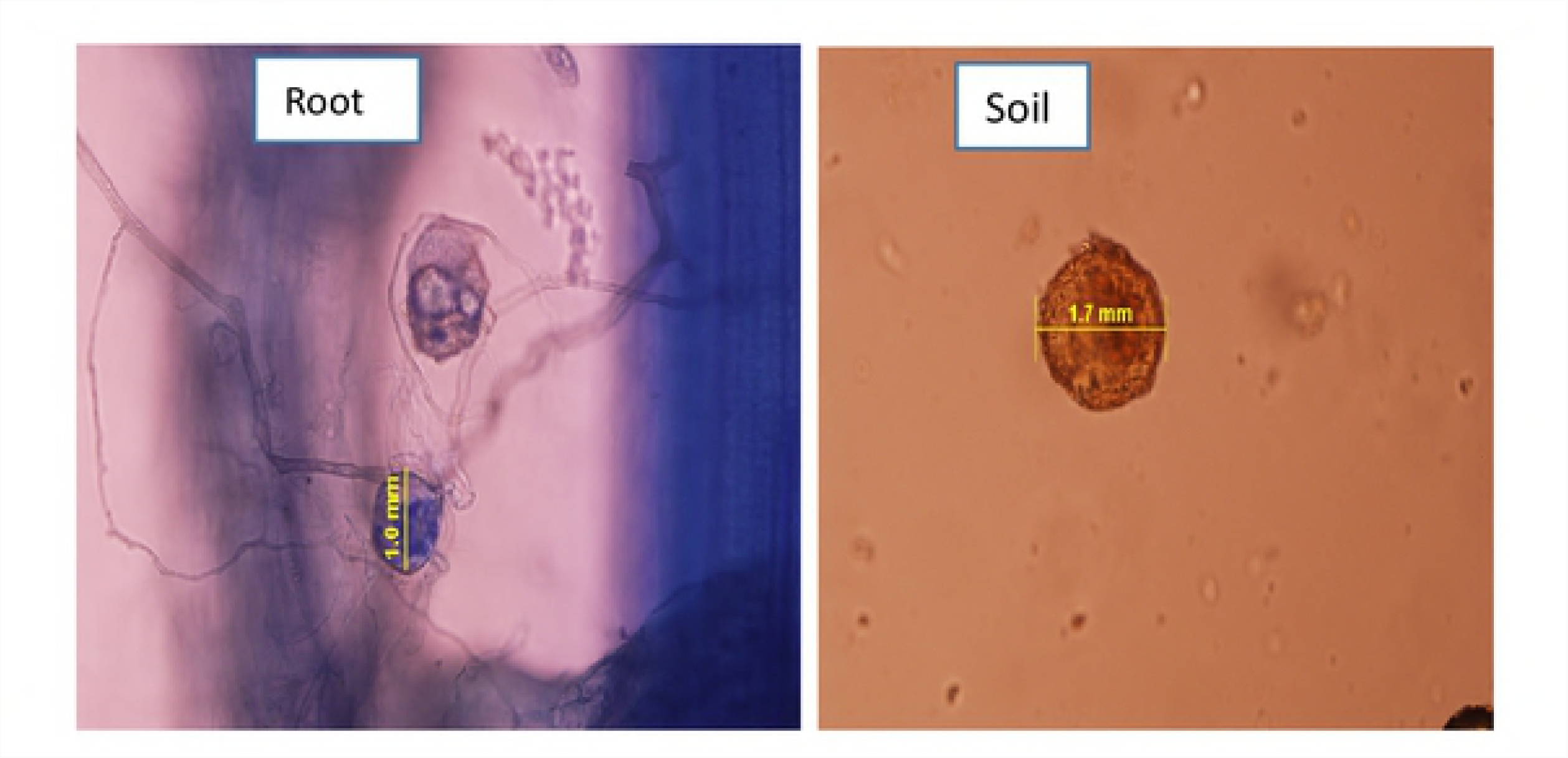
Spore size of AMF in root and soil samples

#### Growing media, green house and sowing time of lentil genotypes

Soils took as growing media for lentil plants in pot experiments. 1200g ground soils kept in each pot for growing lentil. Recommended doses of fertilizers such as, Urea, TSP and MOP were applied to each pot, as in previous experiments. BARI Mashur 1 and BARI Mashur 5 was sown on 13^th^ April 2016 in a controlled temperature greenhouse at BSMRAU. Temperatures ranged from 18°C to 20°C in the greenhouse for lentil growing in pot experiments.

#### Treatments and replications

Two genotypes-BARI Mashur 1 and BARI Mashur 5 were selected and treatments were T_1_ = 8 mgkg^-1^ arsenic concentration in soils, and T_2_ = 45 mgkg^-1^ arsenic concentrated soils. For T_2,_ arsenic concentration increased from 8 mgkg^-1^ to 45 mgkg^-1^ from the source of sodium arsenite (AsNaO_2_). A 150 g of soil with root mixture as Arbuscular Mycorrhizal Fungus (AMF) used for the mitigation of arsenic. Five replications were followed in both AMF and non-AMF treated soils. This experiment was produced total 40 pots for AMF and non-AMF applied soils.

#### Shoot length, fresh and dry weight of root and shoot

Randomly, average shoot length measured through measuring tape (cm) at week 4 in each treated pots. During this week, five lentil plants harvested from arsenic treated each pot. Average fresh weight of root and shoot taken separately through an electrical balance (g) in AMF and non-AMF treated experiment. Similarly, average dry weight of root and shoot of lentil plants measured independently during this week. All samples were dried in an oven at 55°C for 72 hours towards the digestion of samples for the determination of total As accumulation in root and shoot of lentil crops from AMF and non AMF soils.

### 2.4. Digestion of samples

Soils, lentil roots, shoots and pods were digested separately following heating block digestion procedure (Rahman et al., 2007). Of the soil/compost samples, 0.2 g taken into clean, dry digestion tubes and 5 ml of concentrate HNO_3_ and 3 ml concentrate HCLO_4_ added to it. The mixture was allowed to stand overnight under fume hood. In the following day, this vessel put into digestion block for 4 hours at 120° C temperature. Similarly, 0.2 g ground root, shoot and pod samples put into clean digestion vessel and 5 ml concentrate HNO_3_ added to it. The mixture was allowed to stand overnight under fume hood. In the following day, this vessel put into digestion block for 1 hours at 120° C temperature. This content cooled and 3 ml HCLO_4_ added to it. Again, samples put into the heating block for 3-4 hours at 140°C. Generally heating stopped whenever a white dense fume of HCLO_4_ emitted into air. Then samples cooled, diluted to 25ml with de-ionized water and filtered through Whiteman No 42 filter paper for soil and plant samples. Finally, samples were stored with polyethylene bottles. Prior to samples digestion, all glassware was washed with 2% HNO3 followed by rinsing with de-ionized water and drying.

### 2.5. Analysis of total arsenic

Digested samples were brought into the laboratory of Bangladesh Council of Scientific and Industrial Research (BCSIR) for the analysis of total As in lentil root, shoot, pod, soil, vermi-compost and irrigation water. The total As in root, shoot, pod of lentil plants, soil, vermi-compost and water samples were analyzed by flow injection hydride generation atomic absorption spectrophotometry (FI-HG-AAS, Perkin Elmer A Analyst 400) using external calibration (Welsch et al., 1990). The optimum HCl concentration was 10% v/v and 0.4% NaBH_4_ produced the maximum sensitivity. Three replicates taken from each digested samples and the mean values obtained based on the calculation of those three replicates. Standard Reference Materials (SRM) from National Institute of Standards and Technology (NIST), USA analyzed in the same procedure at the start, during and at the end of the measurements to ensure continued accuracy.

### 2.6. Statistical Analysis

The design of this experiment was followed Completely Randomized Block (CRD). Analysis of Variance (ANOVA), means comparison of treatment, varieties, interaction between treatment and varieties, treatment and soils, varieties and soils, treatment-varieties and soils on arsenic accumulation in lentil roots, shoots and pods were analyzed using software R.

## 3. RESULTS

### 3.1 Chemical properties of lentil seed, soil and water samples

The ranges of dry weight of lentil seeds were 9.43 to 9.53 g of 10g BARI released lentil genotypes. Among all lentil cultivars, BARI Mashur 1, BARI Mashur 5, BARI Mashur 6, and BARI Mashur 7 seeds were found As free. The highest As concentration (0.05mgkg^-1^) was found in the seeds of BARI Mashur 4. The distilled water was As free as well as 0.02 mgL^-1^ concentrated arsenic were present in irrigation water. The ranges of pH found 6.75 to 7.93 in vermi-compost, BSMRAU, BJRI and Mathchar soils. The total nitrogen, available phosphorus, exchangeable potassium, and available sulfur were detected 1.23%, 57.71, 150 and 698.04 mgkg^-1^ in vermi-compost samples, accordingly. As well, the total nitrogen, available phosphorus, exchangeable potassium, and available sulfur were detected 0.057%, 14.41, 120 and 9.615 mgkg^-1^ in BJRI soil samples, separately. Similarly, in BSMRAU soil samples, the total nitrogen, available phosphorus, exchangeable potassium, and available sulfur were found 0.11%, 20.68, 124 and 23.07 mgkg^-1^, respectively. On the other hand, Mathchar soil samples content 0.086% of total nitrogen, 9.177 mgkg^-1^ available phosphorus, 128 mgkg^-1^ exchangeable potassium, and 2.884 mgkg^-1^ available sulfur. Total As concentration found 2.688, 8.299, 5.223, and 14.633 mgkg^-1^ in vermi-compost, BJRI, BSMRAU, and Mathchar soil samples, respectively (Table 1).

### 3.2 Biomass and arsenic accumulation in root, shoot and pod of lentil genotypes Shoot length, fresh weight and dry weight of root and shoot of lentil varieties

The highest average shoot length of BARI Mashur 2, BARI Mashur 2 & 3, and BARI Mashur 3 were found 12.5, 11.4, and 9.8 (cm) in T_1_, T_2_ and T_3_ treated lentil seedlings at week 3. T_3_ treated shoot length of BARI Mashur 6 lentil were found significantly lower (*p< 0.001)* than other lentil seedlings (Figure 2). The fresh weight (0.182-0.20 g) was not significantly increased (*p< 0.001)* in T_3_ treated Lentil seedlings. The lowest fresh weight 0.189g was found in T_3_ treated BARI Mashur 5 lentil seedlings at week 3 (Figure 3). In week 9, the ranges of dry weight of root and shoot was found 0.4384 to 0.9064 (g) in As treated lentil seedlings. The highest dry weight of root and shoot were 0.8612 (g) found in BARI Mashur 1 in T_2_ treated seedlings. The lowest was 0.4154 (g) in BARI Mashur 4 of T_2_ treated seedlings. Similarly, the ranges of dry weight of root and shoot were 0.112 to 0.234 (g) in T_3_ treated seedlings. Dry weight of root and shoot were recorded comparatively lower in T_3_ treated lentil seedling than T_1_ and T_2_. Dry weight of root in T_3_ treated BARI Mashur 5 lentil genotypes were found significantly different. As well, Dry weight of shoot in BARI Mashur 7 were found significantly higher than BARI Mashur 1, 2, 4, 5 and 6 lentil genotypes at week 9 (Figures 4, and 5).

**Figure 2.**
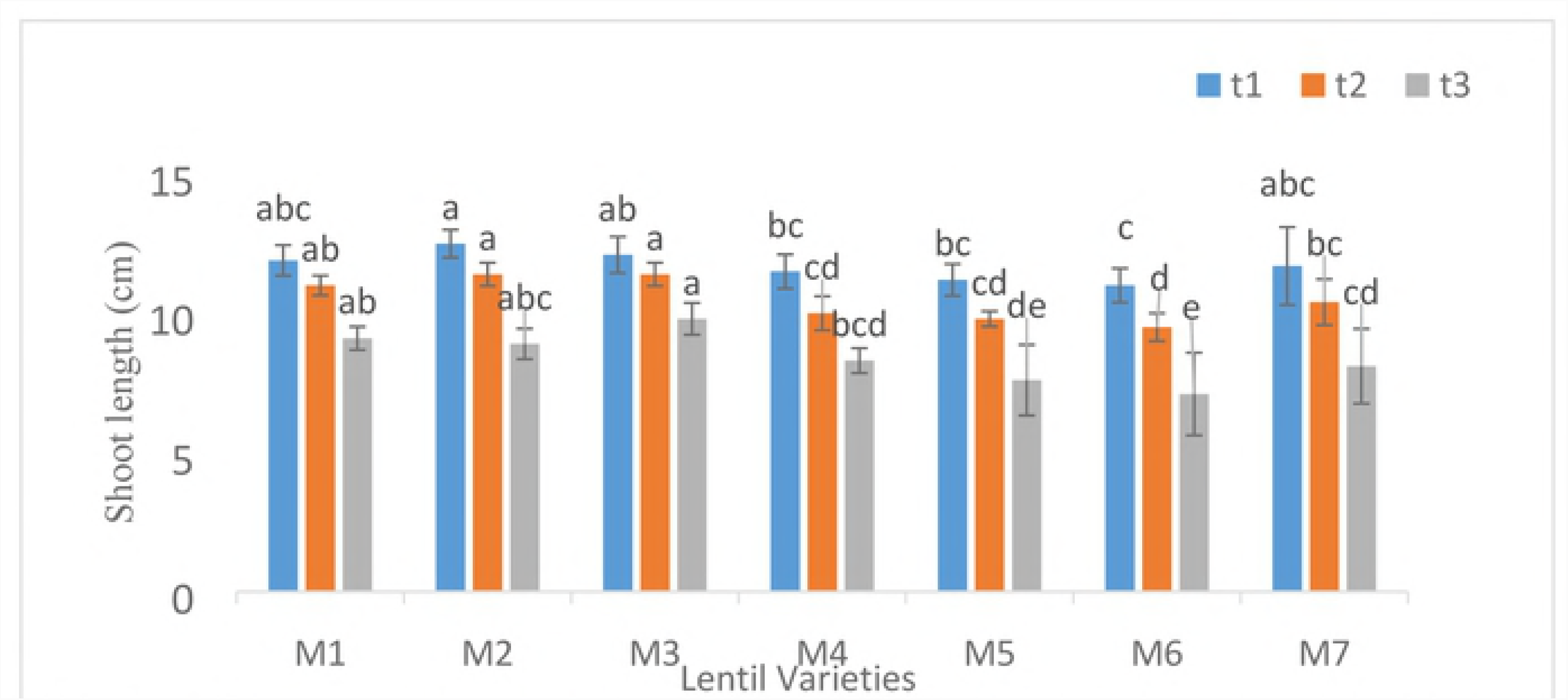
Effect ofarsenic on shoot length (Mean ± SEM) oflentil varieties at week 3. Lettering a b c indicate significantly different at 0.1% level of significance. M1, M2, M3, M4, MS, M6 and M7 indicate BARI Mashur 1, 2, 3, 4, 5, 6, & 7, respectively

#### Arsenic uptake in root and shoot of lentil varieties

According to ANOVA, treatments on arsenic accumulation in root and shoot were found statistically significant (*p< 0.001).* Varieties, and interaction of varieties and treatments both were significantly different on As uptake in lentil roots (*p< 0.001)* (Table 2). Mean comparison of treatment 1 & 2 (*0.001≤p<0.01*), 1 & 3(*p< 0.001),* and 2 &3 (*p< 0.001)* for As accumulation in roots were found significantly difference. As well, the mean comparison of treatment 1 & 3 and treatment 2 & 3 both were found significantly identical (*p< 0.001*) on As accumulation in lentil shoot (Table 3). Interaction of BARI Mashur 1&3 (*0.01≤p<0.0.05*), BARI Mashur1 & 4 (0.05≤p<0.0.1), BARI Mashur1 & 5(*0.001≤p<0.01*), BARI Mashur1 & 6 (0.01≤p<0.05), BARI Mashur 2 & 3 (*p<0.001*), BARI Mashur 2 & 4 (*0.001≤p<0.01*), BARI Mashur 2 & 5 (*p< 0.001*), BARI Mashur 2 & 6 (*0.001≤p<0.01*), and BARI Mashur 2 & 7 (*0.01≤p<0.05*) were found statistically significant on As accumulation in their roots (Table 3). The mean comparison of the interaction between treatments (3) and lentil varieties (7) on As accumulation in root were found statistically significant (*p< 0.001, 0.001≤p<0.01, 0.01≤p< 0.05*) difference (Table 4).

**Table 2.** ANOVA of Arsenic accumulations in root and shoot

| Source of variations (SV) | Degrees of freedom (DF) | Arsenic accumulations in root |  |  | Arsenic accumulations in shoot |  |  |
| --- | --- | --- | --- | --- | --- | --- | --- |
|  |  | Sum of Squares (SS) | Mean Sum of Squares (MSS) | F value | Sum of Squares (SS) | Mean Sum of Squares (MSS) | F value |
| Variety | 6 | 472 | 78.67 | 4.225*** | 151 | 25.17 | 1.607<sup>NS</sup> |
| Treatment | 2 | 18870 | 9435 | 507.001*** | 12647 | 6323.5 | 404.976*** |
| Variety : Treatment | 12 | 756 | 63 | 3.387*** | 303 | 25.25 | 1.617<sup>NS</sup> |
| Residuals | 84 | 1563 | 18.607 |  | 1312 | 15.619 |  |

\*\*\* indicates significant difference at p<0.001 level of significance, <sup>NS</sup> indicates no significant difference

**Table 3.**
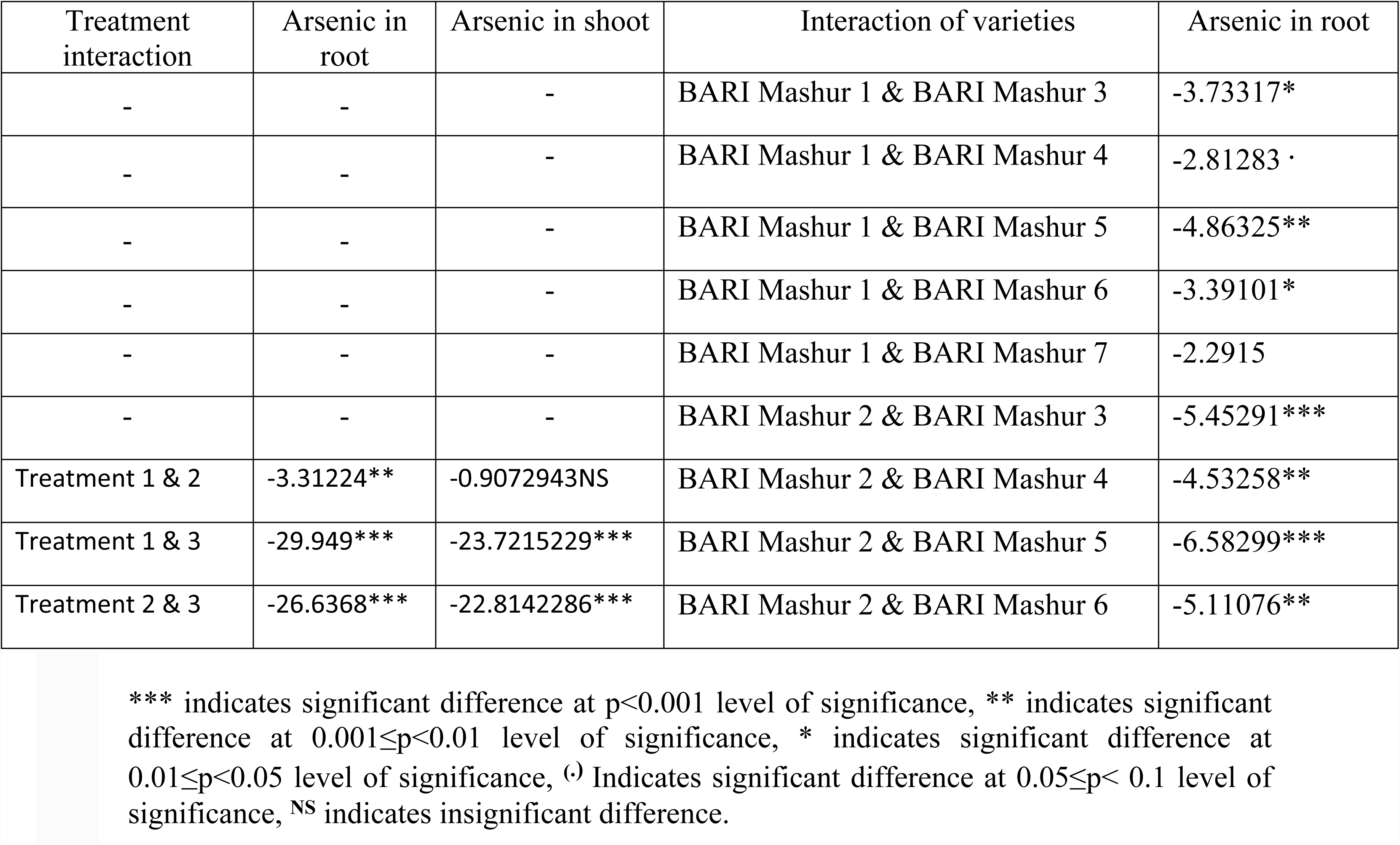
Mean comparison of arsenic accumulations in root and shoot according to the treatment and varieties

**Table 4.** Arsenic accumulation in root according to the interaction between treatment and varieties mean differences

| Comparison (Treatment: Variety -Treatment: Variety) | Arsenic in root |
| --- | --- |
| 3:1-1:1 | 24.7691*** |
| 3:2-1:1 | 18.8576*** |
| 3:3-1:1 | 35.60782*** |
| 3:4-1:1 | 30.74864*** |
| 3:5-1:1 | 36.92746*** |
| 3:6-1:1 | 32.77742*** |
| 3:7-1:1 | 29.89708*** |
| 3:1-2:1 | 22.88976*** |
| 3:2-2:1 | 16.97826*** |
| 3:3-2:1 | 33.72848*** |
| 3:4-2:1 | 28.8693*** |
| 3:5-2:1 | 35.04812*** |
| 3:6-2:1 | 30.89808*** |
| 3:7-2:1 | 28.01774*** |
| 1:2-3:1 | -24.7777*** |
| 2:2-3:1 | -22.1289*** |
| 1:3-3:1 | -24.9863*** |
| 2:3-3:1 | -22.3117*** |
| 3:3-3:1 | 10.83872* |
| 1:4-3:1 | -24.8239*** |
| 2:4-3:1 | -20.376*** |
| 1:5-3:1 | -24.8492*** |
| 2:5-3:1 | -20.3782*** |
| 3:5-3:1 | 12.15836** |
| 1:6-3:1 | -24.8359*** |
| 2:6-3:1 | -20.6582*** |
| 1:7-3:1 | -24.3993*** |
| 2:7-3:1 | -21.513*** |
| 3:2-1:2 | 18.86618*** |
| 3:3-1:2 | 35.6164*** |
| 3:4-1:2 | 30.75722*** |
| 3:5-1:2 | 36.93604*** |
| 3:6-1:2 | 32.786*** |
| 3:7-1:2 | 29.90566*** |
| 3:2-2:2 | 16.21742*** |
| 3:3-2:2 | 32.96764*** |
| 3:4-2:2 | 28.10846*** |
| 3:5-2:2 | 34.28728*** |
| 3:6-2:2 | 30.13724*** |
| 3:7-2:2 | 27.2569*** |
| 1:3-3:2 | -19.0748*** |
| 2:3-3:2 | -16.4002*** |
| 3:3-3:2 | 16.75022*** |
| 1:4-3:2 | -18.9124*** |
| 2:4-3:2 | -14.4645*** |
| 3:4-3:2 | 11.89104** |
| 1:5-3:2 | -18.9377*** |
| 2:5-3:2 | -14.4667*** |
| 3:5-3:2 | 18.06986*** |
| 1:6-3:2 | -18.9244*** |
| 2:6-3:2 | -14.7467*** |
| 3:6-3:2 | 13.91982** |
| 1:7-3:2 | -18.4878*** |
| 2:7-3:2 | -15.6015*** |
| 3:7-3:2 | 11.03948* |
| 3:3-1:3 | 35.82506*** |
| 3:4-1:3 | 30.96588*** |
| 3:5-1:3 | 37.1447*** |
| 3:6-1:3 | 32.99466*** |
| 3:7-1:3 | 30.11432*** |
| 3:3-2:3 | 33.15046*** |
| 3:4-2:3 | 28.29128*** |
| 3:5-2:3 | 34.4701*** |
| 3:6-2:3 | 30.32006*** |
| 3:7-2:3 | 27.43972*** |
| 1:4-3:3 | -35.6627*** |
| 2:4-3:3 | -31.2147*** |
| 1:5-3:3 | -35.688*** |
| 2:5-3:3 | -31.217*** |
| 1:6-3:3 | -35.6746*** |
| 2:6-3:3 | -31.4969*** |
| 1:7-3:3 | -35.238*** |
| 2:7-3:3 | -32.3517*** |
| 3:4-1:4 | 30.80348*** |
| 3:5-1:4 | 36.9823*** |
| 3:6-1:4 | 32.83226*** |
| 3:7-1:4 | 29.95192*** |
| 3:4-2:4 | 26.3555*** |
| 3:5-2:4 | 32.53432*** |
| 3:6-2:4 | 28.38428*** |
| 3:7-2:4 | 25.50394*** |
| 1:5-3:4 | -30.8288*** |
| 2:5-3:4 | -26.3578*** |
| 1:6-3:4 | -30.8155*** |
| 2:6-3:4 | -26.6378*** |
| 1:7-3:4 | -30.3789*** |
| 2:7-3:4 | -27.4926*** |
| 3:5-1:5 | 37.0076*** |
| 3:6-1:5 | 32.85756*** |
| 3:7-1:5 | 29.97722*** |
| 3:5-2:5 | 32.5366*** |
| 3:6-2:5 | 28.38656*** |
| 3:7-2:5 | 25.50622*** |
| 1:6-3:5 | -36.9943*** |
| 2:6-3:5 | -32.8166*** |
| 1:7-3:5 | -36.5577*** |
| 2:7-3:5 | -33.6714*** |
| 3:6-1:6 | 32.84424*** |
| 3:7-1:6 | 29.9639*** |
| 3:6-2:6 | 28.66654*** |
| 3:7-2:6 | 25.7862*** |
| 1:7-3:6 | -32.4076*** |
| 2:7-3:6 | -29.5213*** |
| 3:7-1:7 | 29.5273*** |
| 3:7-2:7 | 26.641*** |

\*\*\* indicates significant difference at $p < 0.001$ level of significance, \*\* indicates significant difference at $0.001 \leq p < 0.01$ level of significance, \* indicates significant difference at $0.01 \leq p < 0.05$ level of significance. For treatment, 1= $T_1$ , 2= $T_2$ and 3= $T_3$ and for variety, 1= BARI Mashur 1, 2= BARI Mashur 2, 3= BARI Mashur 3, 4= BARI Mashur 4, 5= BARI Mashur 5, 6= BARI Mashur 6, and 7= BARI Mashur 7.

#### Arsenic accumulation in pod of lentil varieties during field condition

The collected of BARI released seven lentil varieties were cultivated in 5 mg/kg As concentrated field soils. Among these varieties, BARI Mashur 1 was the highest arsenic (0.45 mgkg^-1^) accumulator and the lowest As (0.029 mgkg^-1^) accumulator was BARI Mashur 7 in its pod. An average As concentration found 0.237, 0.133, 0.298, 0.17, and 0.262 mgkg^-1^ in pods of BARI Mashur 2, BARI Mashur 3, BARI Mashur 4, BARI Mashur 5, and BARI Mashur 6, respectively. Arsenic was significantly increased in pods of BARI Mashur 1 lentil in compare to other genotypes (Figure 6).

### 3.3 Mitigation of arsenic uptake in root and shoot of lentil

#### Spore size of Arbuscular Mycorrhizal Fungus (AMF) in roots and soils

The spore, mycelia and hyphae of AMF observed through stereomicroscope in soil and root samples separately. Sizes of spores were 1-1.7 mm in root samples. On the other hand, spore size of AMF 1.3-1.7 mm was in soil samples. Spore colonization was found 70% in root samples. Number of spore was detected 140 of each kg soil sample (Figure 1).

#### Biomass of lentil genotypes at non-AMF and AMF applied soils

In Non-AMF soils, shoot length of BARI Mashur 1 and BARI Mashur 5 were 6.8 and 6.2 cm in T_1_ treated lentil seedlings. Shoot, length was 5.8, and 3.8 cm were in BARI Mashur 1, and BARI Mashur 5 at T_2_ treated seedlings. AMF treated shoot length at 8 mgkg^-1^ and 45 mgkg^-1^ arsenic concentrated both soils were found significantly higher than non AMF soils during week 4 (Figure 7). Fresh and dry weight of shoot both were found significantly lower in non-AMF treated 45 mgkg^-1^ arsenic concentrated soils at week 5 (Figure 8 and 9). As well as, AMF has significant effect for the increasing of dry and fresh weight of roots in lentil genotypes (Figure 10 and 11).

#### Reduction of arsenic uptake in root and shoot of lentil genotypes

**According to ANOVA,** arsenic accumulation in root and shoot of BARI Mashur 1 and BARI Mashur 5 lentils at non-AMF soils were found significantly difference (p<0.001). As well as, arsenic uptake is significantly reduced in root and shoot of lentil genotypes at AMF treated soils (Table 5). The interaction between treatment & soils on the reduction of As uptake in lentil root and shoot were found statistically significant (p<0.001) (Table 6). Mean comparison effect of the interaction between treatment T_2_ & AMF soils and T_2_ & non AMF soils on the reduction of arsenic accumulation in root and shoot of BARI Mashur 1 and 5 were found statistically significant (p<0.001) (Table 7).

**Table 5.** ANOVA of Arsenic accumulations in root and shoot of BARI Mashur 1 and 5 at non-AMF and AMF soil

| Source of variations (SV) | Degrees of freedom (DF) | Root of BARI Mashur 1 at non-AMF soil |  |  | Shoot of BARI Mashur 1 at non-AMF soil |  |  |
| --- | --- | --- | --- | --- | --- | --- | --- |
|  |  | Sum of Squares (SS) | Mean Sum of Squares (MSS) | F value | Sum of Squares (SS) | Mean Sum of Squares (MSS) | F value |
| Treatment | 1 | 1790.2 | 1790.2 | 1290.23*** | 108.02 | 108.02 | 773.8*** |
| Residuals | 8 | 11.1 | 1.3875 |  | 1.12 | 0.14 |  |
|  |  | Root of BARI Mashur 1 at AMF soil |  |  | Shoot of BARI Mashur 1 at AMF soil |  |  |
| Treatment | 1 | 1070.7 | 1070.7 | 418.2*** | 50.25 | 50.25 | 302.26 *** |
| Residuals | 8 | 20.5 | 2.5625 |  | 1.33 | 0.16625 |  |
|  |  | Root of BARI Mashur 5 at non-AMF soil |  |  | Shoot of BARI Mashur 5 at Non-AMF soil |  |  |
| Treatment | 1 | 745.3 | 745.3 | 641.12*** | 318.7 | 318.7 | 1019.84*** |
| Residuals | 8 | 9.3 | 1.1625 |  | 2.5 | 0.3125 |  |
|  |  | Root of BARI Mashur 5 at AMF soil |  |  | Shoot of BARI Mashur 5 at AMF soil |  |  |
| Treatment | 1 | 392.6 | 392.6 | 640.98*** | 108.25 | 108.25 | 848.24*** |
| Residuals | 8 | 4.9 | 0.6125 |  | 1.7 | 0.2125 |  |

\*\*\* indicates significant difference at p<0.001 level of significance

**Table 6.** ANOVA of arsenic accumulation in root and shoot of BARI Mashur1 and 5 for both soils

| Source of variations (SV) | Degrees of freedom (DF) | Root of BARI Mashur 1 |  |  | Shoot of BARI Mashur 1 |  |  |
| --- | --- | --- | --- | --- | --- | --- | --- |
|  |  | Sum of Squares (SS) | Mean Sum of Squares (MSS) | F value | Sum of Squares (SS) | Mean Sum of Squares (MSS) | F value |
| Treatment | 1 | 2815.0 | 2815.0 | 1428.01*** | 152.81050 | 152.81050 | 999.04*** |
| Soil | 1 | 75.6 | 75.6 | 38.33*** | 8.96594 | 8.96594 | 58.62*** |
| Treat: Soil | 1 | 46.0 | 46.0 | 23.32*** | 5.46117 | 5.46117 | 35.70*** |
| Residuals | 16 | 31.5 | 1.96875 |  | 2.45 | 0.153125 |  |
|  |  | Root of BARI Mashur 5 |  |  | Shoot of BARI Mashur 5 |  |  |
| Treatment | 1 | 1109.8 | 1109.8 | 1250.48*** | 399.2 | 399.2 | 1520.76*** |
| Soil | 1 | 53.9 | 53.9 | 60.73*** | 44.2 | 44.2 | 168.38*** |
| Treat: Soil | 1 | 28.0 | 28.0 | 31.55*** | 27.7 | 27.7 | 105.52*** |
| Residuals | 16 | 14.2 | 0.8875 |  | 4.2 | 0.2625 |  |

\*\*\* indicates significant difference at p<0.001 level of significance

**Table 7.** Mean comparison of the interaction between treatment and soils on the reduction of arsenic accumulation in root and shoot of BARI Mashur 1 and 5 for both soils

| Comparison | Arsenic in root of BARI Mashur 1 | Arsenic in shoot of BARI Mashur 1 | Arsenic in root of BARI Mashur 5 | Arsenic in shoot of BARI Mashur 5 |
| --- | --- | --- | --- | --- |
| T<sub>2</sub>: non AMF - T<sub>1</sub>: non AMF | 26.7598*** | 6.5734*** | 17.266*** | 11.29*** |
| T<sub>1</sub>: AMF - T<sub>1</sub>: non AMF | -0.8552<sup>NS</sup> | -0.294<sup>NS</sup> | -0.9152<sup>NS</sup> | -0.6176<sup>NS</sup> |
| T<sub>2</sub>: AMF - T<sub>1</sub>: non AMF | 19.84*** | 4.1892*** | 11.616*** | 5.9626*** |
| T<sub>1</sub>: AMF - T<sub>2</sub>: non AMF | -27.615*** | -6.8674*** | -18.1812*** | -11.9076*** |
| T<sub>2</sub>: AMF - T<sub>2</sub>: non AMF | -6.9198*** | -2.3842*** | -5.65*** | -5.3274*** |
| T<sub>2</sub>: AMF - T<sub>1</sub>: AMF | 20.6952*** | 4.4832*** | 12.5312*** | 6.5802*** |

\*\*\* indicates significant difference at p<0.001 level of significance, <sup>NS</sup> indicates insignificant difference

Treatment, variety, and treatment & varietal interaction effect in root and shoot at AMF and non AMF soil were found statistically significant (p<0.001) (Table 8). Mean comparison effect of the interaction between T_2_ & BARI Mashur 1 and T_1_ & BARI Mashur 1; T_2_ & BARI Mashur 5 and T_1_ & BARI Mashur 1; T_1_ & BARI Mashur 5 and T_2_ & BARI Mashur 1; T_2_ & BARI Mashur 5 and T_2_ & BARI Mashur 1; and T_2_ & BARI Mashur 5 and T_1_ & BARI Mashur 5 (p<0.001) were found statistically significant difference on As accumulation in their root and shoot at non-AMF soils. As well as, in AMF soils, arsenic accumulation was significantly reduced (p<0.001) in their root and shoot of both lentil varieties (Table 9).

**Table 8.** ANOVA of arsenic accumulation in root and shoot according to treatment and varieties for non-AMF and AMF soil

| Source of variations (SV) | Degrees of freedom (DF) | Root at non-AMF soil |  |  | Shoot at non-AMF soil |  |  |
| --- | --- | --- | --- | --- | --- | --- | --- |
|  |  | Sum of Squares (SS) | Mean Sum of Squares (MSS) | F value | Sum of Squares (SS) | Mean Sum of Squares (MSS) | F value |
| Treatment | 1 | 2422.8 | 2422.8 | 1909.596*** | 398.9 | 398.9 | 1772.89*** |
| Variety | 1 | 120.2 | 120.2 | 94.739*** | 53.2 | 53.2 | 236.44*** |
| Treat: Variety | 1 | 112.7 | 112.7 | 88.828*** | 27.8 | 27.8 | 123.56*** |
| Residuals | 16 | 20.3 | 1.26875 |  | 3.6 | 0.225 |  |
|  |  | Root at AMF soil |  |  | Shoot at AMF soil |  |  |
| Treatment | 1 | 1380.0 | 1380.0 | 869.29*** | 153.00 | 153.00 | 807.92*** |
| Variety | 1 | 92.4 | 92.4 | 58.21*** | 13.25 | 13.25 | 69.97*** |
| Treat: Variety | 1 | 83.3 | 83.3 | 52.47*** | 5.50 | 5.50 | 29.04*** |
| Residuals | 16 | 25.4 | 1.5875 |  | 3.03 | 0.189375 |  |

\*\*\* indicate significant difference at p<0.001 level of significance

**Table 9.**
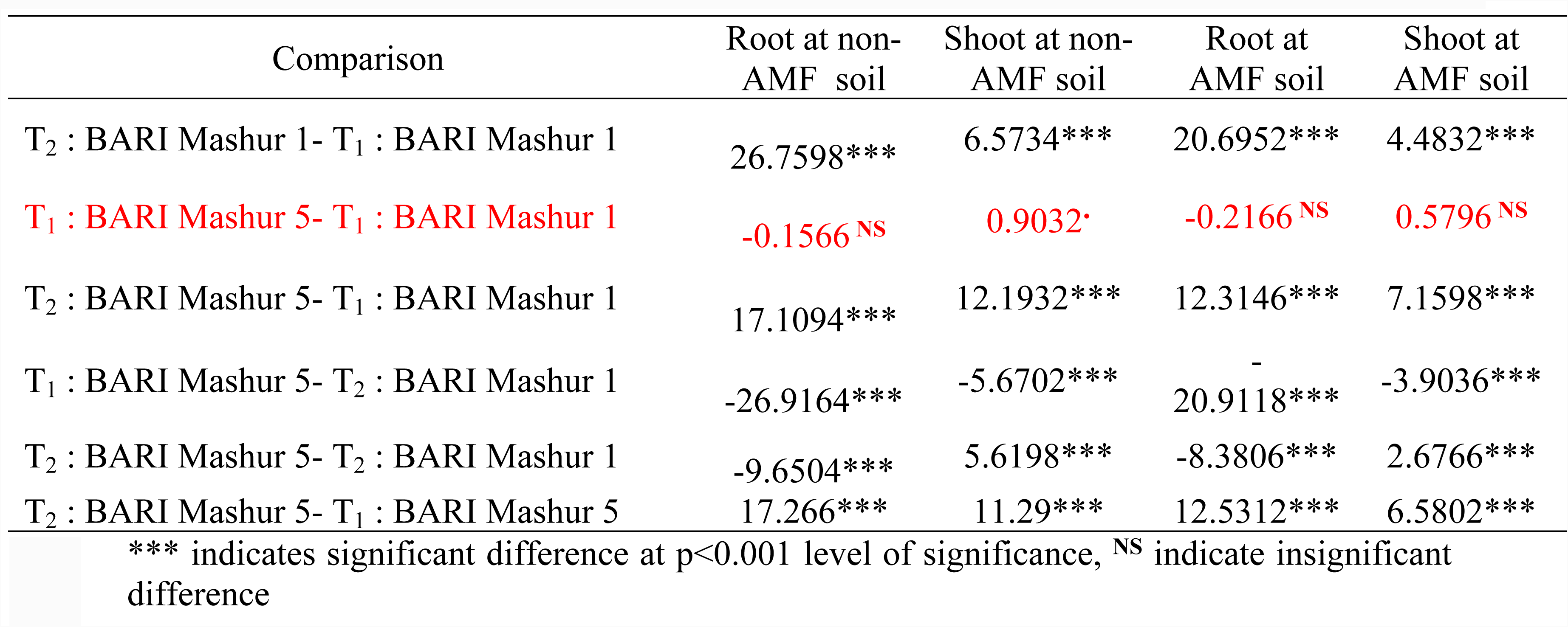
Means comparison of an interaction effect between treatment and varieties on arsenic accumulations in root and shoot for non-AMF and AMF soils.

According to ANOVA, treatment, variety, soil, treatment & varietal interaction, and treatment & soil interaction effect in root and shoot of lentil plants were found statistically significant (p<0.001). On the other hand, the interaction between variety & soil; treatment, variety and soil were found statistically significant difference (p<0.001) in shoot (Table 10). According to the interaction between treatment and soils, mean comparison effect of the interaction between T_2_ & AMF and T_2_ & non-AMF soils on the reduction of As uptake in root and shoot of both lentil crops were found statistically significant (p<0.001) (Table 11). According to the interaction between variety and soils, means comparison of the interaction effect of BARI Mashur 5& AMF and BARI Mashur 5 & non-AMF soils on the reduction of As uptake in lentil shoot in this pot experiment were found to be statistically significant p<0.001) (Table 11). Also, interaction effect between treatment, variety and soil, on the reduction of As uptake in shoot of lentil crops were found statistically significant difference (p<0.001; 0.001≤p<0.01) (Table 12).

**Table 10.** ANOVA of arsenic accumulation in root and shoot according to treatment, varieties and soils in pot experiment

| Source of variations (SV) | Degrees of freedom (DF) | Arsenic accumulations in root |  |  | Arsenic accumulations in shoot |  |  |
| --- | --- | --- | --- | --- | --- | --- | --- |
|  |  | Sum of Squares (SS) | Mean Sum of Squares (MSS) | F value | Sum of Squares (SS) | Mean Sum of Squares (MSS) | F value |
| Treatment | 1 | 3730 | 3730 | 2594.78*** | 523.0 | 523.0 | 2497.91*** |
| Variety | 1 | 212 | 212 | 147.48*** | 59.8 | 59.8 | 285.61*** |
| Soil | 1 | 129 | 129 | 89.74*** | 46.5 | 46.5 | 222.09*** |
| Treat: Variety | 1 | 195 | 195 | 135.65*** | 29.0 | 29.0 | 138.51*** |
| Treat: Soil | 1 | 73 | 73 | 50.78*** | 28.9 | 28.9 | 138.03*** |
| Variety : Soil | 1 | 1 | 1 | 0.696<sup>NS</sup> | 6.7 | 6.7 | 32.02*** |
| T:V:S | 1 | 1 | 1 | 0.696<sup>NS</sup> | 4.3 | 4.3 | 20.54*** |
| Residuals | 32 | 46 | 1.4375 |  | 6.7 | 0.209375 |  |

\*\*\* indicates significant difference at p<0.001 level of significance, <sup>NS</sup> indicate insignificant difference

**Table 11.**
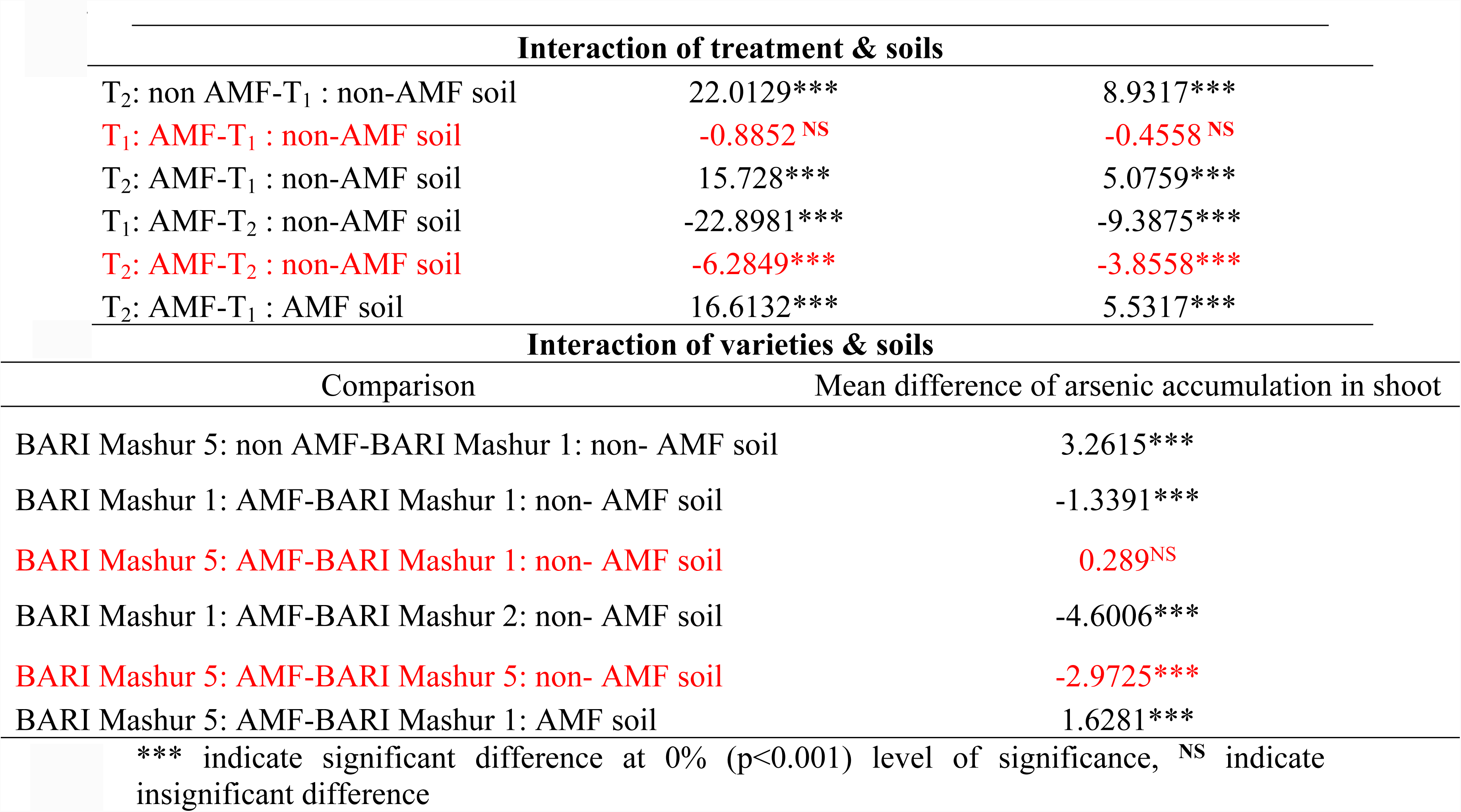
Mean comparison of arsenic accumulation in root and shoot for both lentil varieties according to the interaction of treatment & soils, and varieties & soils in pot experiment

**Table 12.**
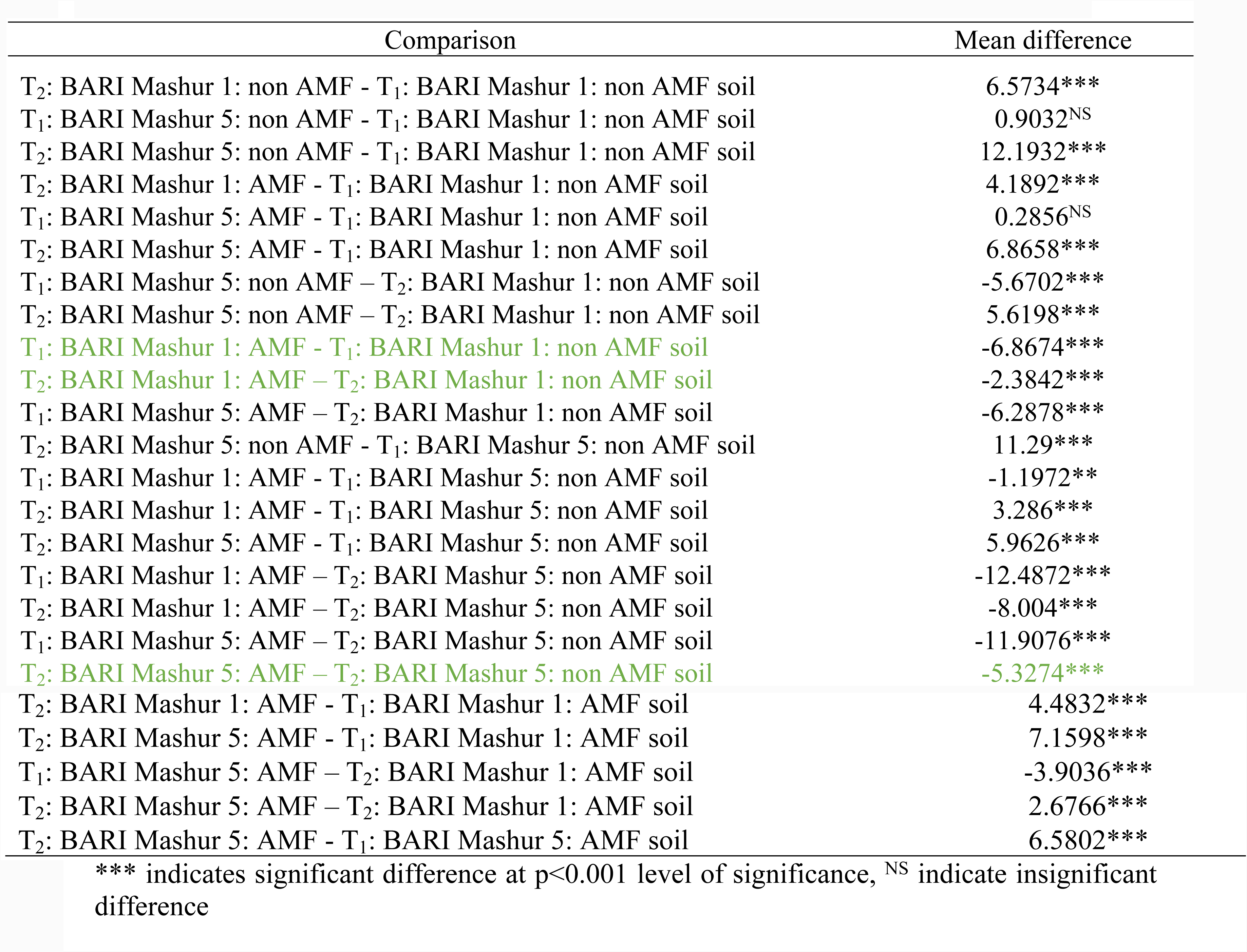
Means comparison of an interaction effect between treatment, varieties and soils on arsenic accumulations in shoot

## 3. Discussion

Arsenic (As) contamination in soils has been reported in many countries throughout the world, with the most severe problems found in Asia, particularly Bangladesh (Chowdhury et al., 1999; Dhar et al., 1997). In Bangladesh, the contamination of As in groundwater was confirmed in 1993 (Tondel et al., 1999). Since then, this contamination has been extended to crop fields due to the irrigation of ground water in Bangladesh (Alam et al., 2011; Tondel et al., 1999). Among several contaminated areas, Faridpur region is one of the highest As contaminated in Bangladesh. Most of these areas are As polluted due to highly uses of ground water irrigation in their crop fields. We found about 15 mg/kg concentrated of arsenic in background soils of these regions, this concentration is definitely dangerous for the development of root, shoot and grains for many cereal crops as well as lentil plants (Table 1). Similarly, As contamination in food crops is also highly visible in other region of Bangladesh including west India (Ullah, 1998; Alam and Sattar, 2000).

Lentil is one of the important leguminous food crops as well as rice and other minor cereal crops in Bangladesh. Plant’s protein is significantly essential for physiological growth of human beings. Nevertheless, these food crops have contaminated because of high concentrated As presence in soils of crop fields. Generally, lentil grown in dry season, so irrigation needed for successful cultivation of this crop. Arsenic in background soils and water lead to elevate the concentration of As in lentil root, shoot and grain (Ahmed et al., 2006). These type of uptake in root, shoot and pod of lentil crops is connected with several nutrient in soils specially phosphate content in soils (Ahmed et al., 2006; Hingston et al., 1972). We found phosphorus concentration (9-57 mgkg^-1^) in soil samples for pot experiment, which increased the As accumulation in lentil root, shoot and pods (Table 1).

Arsenic accumulation in lentil genotypes has significantly affected on its biomass. Different vegetative responses of lentil plants such as root length, shoot height, root and shoot biomass had studied in this experiment (Figure 3 and 4). Kapustka et al. (1995) reports the sensitivity of vegetative response follows the order: root length>root mass>shoot length>total mass (root + shoot)>shoot mass>germination. However, we found As sensitivity was higher on lentil’s roots, shoots, and pods, accordingly. Shoot, height, fresh weight, dry weight of root and shoots, plant biomass (root + shoot + pod) and root length were significantly affected with increasing of As concentration in soils. For instance, total biomass of lentil crops was found to be in more jeopardy in 100 mgkg^-1^ As concentrated soils than other treated pots (5 mgkg^-1^ As; 15 mg kg^-1^ As) of lentil seedlings (Figures 3, 4, and 5).

BARI released all lentil are promising varieties in Bangladesh as well as throughout the world. Not yet conducted of an experiment against As uptake from soil to root, shoot and grain in lentil of Bangladesh. In fact, Bangladesh is the second largest As contaminated region throughout the world. In addition, lentil is the number one pulse crops as a source of protein. Humans need more protein for the proper development of their immune system. In this regards, lentil is also one of the cheapest sources of protein for the effort on mental development. This protein should be toxin free and healthy to consume for human beings. However, all lentil varieties were performed with significant differences for the accumulation of As in their roots in 5, 15 and 100 mgkg^-1^ concentrated soils due to the less genotypic variation. Nevertheless, As accumulation is not significantly increased in shoots and pods of lentil plants. We found all lentil varieties were grown in good condition during seedling stage in 5 and 15 mgkg^-1^ arsenic concentrated soils compare to the 100mgkg^-1^ concentrated soils (Figures 3, 4, and 5).

This is good news that not significant concentration of As has transported from soils to lentil pods (Table 5). In fact, BARI Mashur 1 genotypes were identified higher As accumulator (0.45 mgkg^-1^) in pods than other genotypes (Figure 6). Similarly, irrespective of As dose, roots contained higher concentration of As than shoots and pods. Higher As concentration in roots reported by Marin et al. (1992, 1993), Xie and Huang (1998) and Abedin et al. (2002) in food crops. There are, however, no previous reports of elevated As concentrations in lentil pods. This research has significant importance in terms of human food chain. Lentil pods, root and shoot are highly used as food for humans, and animals throughout the world. Arsenic might have been transferred to human bodies through the food chains. This transportation is conditional on the availability of As in soils from its source. It has carcinogenic effect in the Bengal Delta Plain is considered to be the largest mass poisoning in the history of humanity as millions of people are exposed and suffer the effects of chronic As intoxication (Smith et al., 2008). Arsenic has identified as a non-threshold human carcinogen (International Agency for Cancer Research [IARC], 2004). Furthermore, other than cancer, human exposure to As has been associated to diverse health problems such as cardiovascular disease, skin lesions, and diabetes (World Health Organization [WHO], 2011). The concentration of As in the groundwater in Bangladesh and West Bengal (India) exceeds by several times the permissible levels set internationally and nationally (Chakraborti et al., 2009; Mandal and Suzuki, 2002). Due to the critical situation, arsenic free lentil grains/pods are significantly important in the South Asian network as well as all over the world.

In these circumstances, low As accumulator lentil genotypes are important for human beings. For this mitigation of arsenic, AMF can reduce the As uptake in root, shoot and pods of lentil crops (Orlowska et al., 2012). This AMF colonized with lentil roots, which is deterred As uptake and As toxicity through the symbiosis relationship between each other. It is consistently enhanced the reduction of As toxicity, and plants generally show increases in growth compared with Non-AMF controls grown at the same As and P supplies in soil (Ahmed et al., 2006; Covey et al., 1981; Pope et al., 2007; Ultra et al., 2007b; Xia et al., 2007). We found BARI Mashur 1 and BARI Mashur 5 both lentil genotypes performed better for their growth of root and shoots in 8 mgkg^-1^ and 45 mgkg^-1^ arsenic concentrated AMF applied soils than non-AMF. We also found shoot length, dry weight of shoot and root, fresh weight of root and shoot of lentil were higher in AMF treated soil than non-AMF applied soils. Root and shoot, growth was satisfactory of both varieties of lentil in mutually treated of AMF applied soils (Figure 7-11).

Arsenic has increased significantly in root and shoot of BARI Mashur 1 and BARI Mashur 5 of lentil genotypes. There is also evidence AMF can reduce As uptake in root and shoot in both lentil genotypes (Table 7 and 8). Research also showed that AMF have their substantial effect on plant growth. The growth parameters decreased significantly with the increase rate of As concentration in soils. It emphasized that AMF inoculation reduced As translocation from soil to plant and increase growth and nutrient uptake and chlorophyll content of food crops significantly (Elahi et al., 2010). Similarly, there is growing evidence that Mycorrhizal fungi might alleviate As toxicity to the host plant by acting as a barrier in soils (Leyval et al. 1997). It has been widely reported that mycorrhizas fungi can increase the tolerance of their host plants to heavy metals when present at toxic levels (Bradley et al., 1982; Jones and Hutchinson 1988). Heggo and Angle (1990) and Hetrick et al. (1994) as well as demonstrated that, at high level of As concentration in soils, AMF infection reduced the concentration of As in plant biomass.

Plant growth changes due to the presence of toxic substances and availability of nutrient in soils. Arsenic toxicity is one of the important factors for the nutrient availability in soils, which directly deterred to stunt of plant growth. For this, we need to improve soil health condition through the mitigation process of arsenic toxicity in soils. As a reason, we used AMF for the improvement of soil condition through the mitigation of arsenic toxicity in soils. There is also evidence AMF can be effective in 8 and 45 mgkg^-1^ arsenic concentrated soils for the reduction of arsenic uptake in root and shoot from soils (Table 11 and 12). Similar result also found that AMF play an important role in protecting crop plants against As contamination. However, this is the direct involvement of arbuscular mycorrhizal fungi (AMF) in detoxification mechanisms. AMF treated soils indicate that fungal colonization dramatically increased plant’s biomass growth (Chen et al., 2007). Research demonstrate a positive effect of mycorrhizal inoculation on growth of lentil (*L*. *culinaris*), P nutrition, and lessens As toxicity in plant soil interaction (Chen et al., 2007). It can reduce into human body through food chains using AMF inoculation in As contaminated soils. Reduced uptake of As by lentil roots and subsequently, transformation to shoots and pods, has particularly will not be implicated to the human food chain.

## 4. Conclusion

Arsenic is the number one carcinogenic substances. Among 37 countries, Bangladesh is one of the second largest arsenic contaminated areas in the world. Not only Bangladesh, many countries has identified As is the toxic and hazardous substances. Lentil is one of the important legume crops in Bangladesh as well as throughout the world as a source of protein. This source of protein should have confirmed toxin free for human beings. For this reason, accumulation of As and its mitigation in lentil genotypes is significantly important for the future demand of food safety. We found BARI Mashur 1 lentil genotypes high As accumulator than other released lentil varieties in Bangladesh. AMF applied for the mitigation of As from soils to root, shoot and pods in these lentil genotypes. We found AMF could effectively reduce As transportation from soil to root and shoot of lentil seedlings. It also diagnosed that AMF has decreased As uptake in root and shoot of lentil crops. Therefore, the mitigation of As in lentil root, shoot and pod is significantly important for the supplying of toxin free lentil seeds throughout the world using AMF in soils.

## Acknowledgement

Authors are grateful to the laboratory of Crop and Soil Sciences at **Washington State University, WA, USA for their research support**. The authors also thank to the Laboratory of Environmental Science at BSMRAU, soil science at BAU and Biological Research Division at Soil and Environment Section of BCSIR. Finally, we are especially thankful to the ASPADA for their valuable funding on behalf of this research project.

